# Decoupled seasonal effects of an environmentally transmitted wildlife disease

**DOI:** 10.64898/2026.08.17.743117

**Authors:** Macy J. Kailing, Louise Callanan, Marta Valldeperes, Shane A. Richards, Scott Carver

## Abstract

1. Seasonal forcing is a dominant factor shaping host-pathogen interactions and disease dynamics across many wildlife systems, including species impacted by environmentally transmitted parasites. How seasonality in parasite dynamics translates to the host when the infection and disease impacts operate at different timescales, however, remains poorly understood.
2. We investigate how seasonality shapes sarcoptic mange in bare-nosed wombats, *Vombatus ursinus*, a disease caused by the environmentally transmitted parasitic mite *Sarcoptes scabiei*, causing a protracted clinical time-course in the host. Using an empirically informed state-based deterministic model we explore how wombat population trajectories are influenced by (i) seasonal constraints to mite survival and (ii) in context of host-pathogen encounter rates, as measured by the ratio of burrows to wombats.
3. We demonstrate three long-term outcomes of wombat-mange: host and parasite extinction, endemic disease, and disease-free. We find seasonal environments narrow the range of host-pathogen encounter rates that support *S. scabiei* persistence relative to stable environments, and prevalence and population sizes vary more in seasonal compared to stable environments except under moderate host-pathogen encounter rates when seasonal effects are less apparent. We also find that a protracted infectious period is essential for host-parasite coexistence in the wombat-mange system.
4. Our seasonal model results are consistent with field observations, such that mange prevalence in natural populations increases during seasons of longer off-host mite survival. Application of these findings suggest management efforts could reduce host population impacts through disease management in seasons with longer off-host parasite survival or reduce the environmental reservoir through disease management in seasons with shorter off-host survival.
5. We provide novel, mechanistic explanations for distinctive population trajectories that arise from a seasonally forced wildlife disease, including climate factors that operate independently on parasites, host demography, and disparate timescales over which seasonality affects parasites and hosts. Broadly, linking seasonality to long-term population dynamics can improve the predictability and management of wildlife diseases, but requires an understanding of how local intrinsic factors interact with seasonal pressures over time.

## 1 Introduction

Seasonal factors are a ubiquitous feature of infectious diseases that can force periodic epidemics by altering host-pathogen interactions (Altizer et al. 2006, Martinez-Bakker et al. 2015, Martinez 2018). Across wide-ranging human and animal systems, seasonal fluctuations in climate, species interactions, or resource availability drive the timing of outbreaks by affecting host immune defense, host aggregations and contacts, and birth pulses (Altizer et al. 2004, Hosseini et al. 2004, Altizer et al. 2006, Langwig et al. 2015, Hirsch et al. 2016, Wilber et al. 2022). Despite seasonal drivers of infection being common, how they influence the periodicity of epidemics long term or disease persistence remains poorly understood for many wildlife diseases. Attributing seasonal forcing to long-term outcomes is difficult because it arises through complex processes involving the traits of hosts and pathogens that seasonality is acting on, the seasonal mechanism that is forcing infections, and how the seasonally affected epidemiological parameter (i.e., contact, birth rate, pathogen decay) interacts with intrinsic conditions that vary over time, such as local host density (Altizer et al. 2006, Grassly et al. 2006, Martinez 2018).

Temporal variation in biological processes of hosts and parasites is an additional source of variation in disease dynamics, but the importance of seasonality in this context is less understood. For example, temperature driven pulses in vector abundance can occur over days to weeks while susceptible host recruitment and abundance changes over months or years, resulting in a lagged host population response and chaotic disease dynamics (Koolhof et al. 2024). In addition to demography, the timing of transmission-relevant processes is also predicted to differ between hosts and pathogens and govern long-term dynamics. For example, climate induced changes to vector bite rates can periodically amplify transmission between hosts, but pathogen persistence long term can depend on the duration of the infectious period relative to vector seasonality (Khong et al. 2023). A lack of understanding the importance of temporal variation in host and pathogen demography as well as the timespans of infection and disease relative to seasonal fluctuations presents barriers to describing the causes and consequences of seasonally forced host-pathogen dynamics. While challenging, characterizing seasonal processes that control population responses across a range of host conditions is critical for anticipating impacts and developing management strategies for wildlife diseases.

Pathogens that persist in the environment, in reservoirs or on fomites, cause a uniquely consequential group of diseases, owing to their ability to initiate new epidemics, sustain outbreaks, and facilitate cross-species transmission (De Castro et al. 2004, Turner et al. 2016, Hopkins et al. 2022, Vargas Soto et al. 2026). Notable examples in wildlife include chytrid fungus in amphibians (Mitchell et al. 2007, Kilpatrick et al. 2010), white nose syndrome in bats (Hoyt et al. 2018, Hoyt et al. 2021), chronic wasting disease in cervids (Haley et al. 2015), and sarcoptic mange in a range of wildlife species (Browne et al. 2021, Escobar et al. 2022). Environmental reservoir dynamics – such as the establishment, distribution, or maintenance of infectious areas – are prone to seasonal effects. Environmental fluctuations can influence reservoir dynamics through several mechanisms, such as off-host pathogen survival (Breban et al. 2009, Rohani et al. 2009, Hoyt et al. 2020), host encounter rates (e.g. density-based processes or seasonal movement patterns as examples) (Almberg et al. 2011, Dolfi et al. 2024), or pathogen shedding rate from hosts (King et al. 2015, Laggan et al. 2022). By regulating transmission risk through environmental reservoirs, seasonality can be a key determinant of the long-term outcomes of diseases that are environmentally transmitted (Breban et al. 2009, Rohani et al. 2009, Hoyt et al. 2018, Wilber et al. 2022). However, because estimating susceptible host encounters with contaminated reservoirs and pathogen survival rates jointly across seasons and under variable host contexts is difficult, the links between seasonality, transmission, and host population outcomes remain a critical knowledge gap for many wildlife diseases that are environmentally transmitted.

Sarcoptic mange is the most significant disease affecting bare-nosed wombats, *Vombatus ursinus* (Martin et al. 2018). Caused by the parasitic mite, *S. scabiei* (Yabsley et al. 2025), and introduced to Australia through European colonialism, the parasite has established within bare-nosed wombat populations throughout their southeastern Australian distribution (Fraser et al. 2016, Fraser et al. 2018). A range of epidemiological dynamics are observed, including epidemics with host decline, endemic disease, and disease-free areas (Carver et al. 2023). Evidence supports environmental transmission occurring within the burrow of this fossorial marsupial, owing to: bare-nosed wombats being relatively solitary in lifestyle with few direct contacts outside of mating (Carver et al. 2024); asynchronous burrow sharing through burrow-switching every 1-9 days (Martin et al. 2019); exposure of healthy bare-nosed wombats to mites left behind by a former infested occupant (Martin et al. 2018, Driessen et al. 2021); and little evidence of other species involvement beyond rare cross-species exposures. Wombats infested by *S. scabiei* suffer a protracted period of progressively severe clinical crusted mange signs, including alopecia, hyperkeratosis, emaciation and eventual death, spanning approximately three months (Skerratt 2005, Martin et al. 2018). There is currently little evidence of resistance to *S. scabiei* induced mortality and variable epidemiological outcomes suggest multiple non-immune processes can drive population outcomes (Carver et al. 2023). Importantly, research suggests seasonal forcing in the environmental reservoir stage of *S. scabiei* (Browne et al. 2021), with survival estimated to fluctuate from 5-16 days across summer and winter conditions, respectively (Arlian et al. 1984, Arlian et al. 1989, Browne et al. 2021), although the impact this has on epidemiological dynamics of sarcoptic mange in bare-nosed wombats has not been investigated.

Our overarching aim was to investigate how the parasite in our system (*S. scabiei*) with strong seasonality in environmental survival shapes disease dynamics in the host (wombat), given divergent timescales in the host, relative to the environment. We integrate information on seasonal constraints in environmental mite survival across a range of host conditions – specifically, the ratios of burrows to wombats that influence environmental encounter rates – to understand seasonal context of epidemiological outcomes for wombat-mange. We use an empirically informed state-based mechanistic model to address three objectives, each focused on the interactive effects of seasonality and initial host-pathogen encounter rates: 1) evaluate long-term epidemic outcomes; 2) explore temporal variation in local host density, outbreak dynamics, and reservoir persistence; 3) evaluate thresholds mediated by seasonality in parasite dynamics, and 4) compare the epidemiological outcomes of our model to empirical trends. We show that seasonality in mite environmental survival constrains the range of burrow to wombat ratios over which *S. scabiei* can persist, relative to stable mite survival, and that epidemiological dynamics in the host are influenced by, but decoupled from, seasonal patterns of mite survival in the environment. Our modelling outcomes demonstrate that seasonality in mite environmental survival is both epidemiologically important and produces enigmatic effects broadly consistent with field observations.

## 2 Materials and Methods

### 2.1 Model description, parameter estimation, and simulations

We worked with an established discrete-time state-based model published by Martin et al. (2019), previously shown to capture processes shaping host and pathogen dynamics in the wombat-*S. scabiei* system. In contrast to typical host-disease models that estimate continuous rates, this state-based approach estimates terms as daily probabilities that accommodate the range of biological timescales that are operating in this system, including burrow switching (days), disease progression (months), and wombat reproduction (years). The model framework consists of a matrix of burrow reservoirs where the fraction of burrows existing in one of eight states is calculated at daily timesteps. The eight possible states of burrows are based on the presence or absence of mites as well as wombat presence and infection status (Figure 1A). The state of a burrow can change daily based on five event probabilities in the following sequence, 1) wombat mortality, 2) unoccupied, contaminated burrows lose mites, 3) low infected wombats progress to high infected, 4) wombat switches burrows, which also determines event 6 of Figure 1A, and 5) wombat offspring gains independence. We used the same parameter estimates for the daily state transitions as in Martin et al. (2019) with slight modifications as detailed in the Supplemental Methods. Most distinctly, we transform the probability that unoccupied, contaminated burrows lose mites into a time-varying term to accommodate seasonal effects on mite persistence off host.

**Figure 1.**
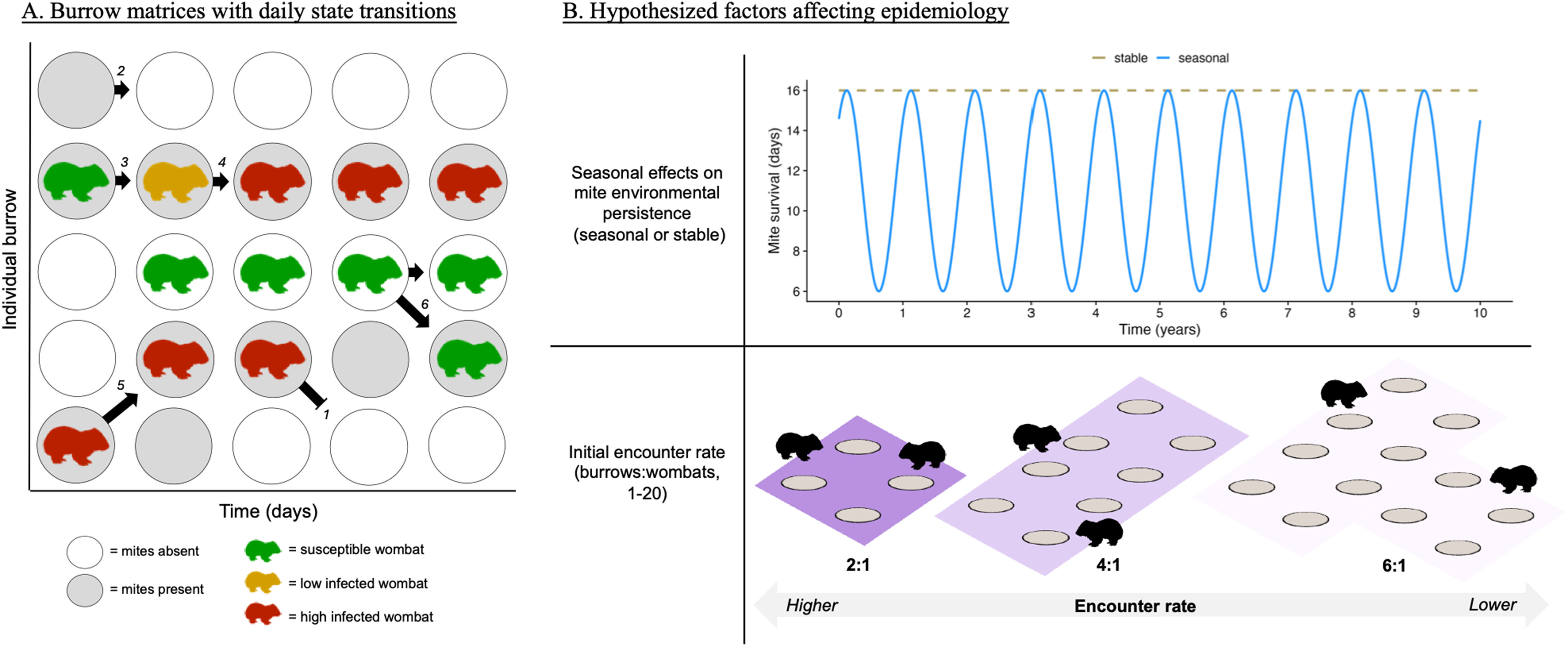
Conceptual diagram of the model framework and the hypotheses tested. **A)** Diagram of the state-based, discrete-time model. Parameters controlling daily state transitions are defined in Supplemental Table 1 with examples shown in the figure as *1* = wombat mortality; *2* = loss of mites from burrow; 3 = Susceptible wombat acquires infection; 4 = disease progression; 5 = wombat switches burrow; 6 = joey gains independence. **B)** *Top*: Mite survival predictions given over time in stable (tan-dashed) and seasonal (blue-solid) environments. *Bottom*: Depiction of the relationship between initial burrows:wombats and encounter rates. Fewer burrows available per wombat (2:1, darkest purple) increases the frequency of burrow sharing while more burrows available to wombats (6:1, lightest purple) decreases burrow sharing. The relative encounter rate decreases as the burrows:wombats ratio increases. Note that the range of initial burrows:wombats explored in simulations extended from 1 to 20 but also varied between time steps as the fraction of burrows occupied (n, relative host density) changed.

We incorporated seasonality to the model based on temperature and humidity dependent mite survival estimates in wombat burrows from (Browne et al. 2021). To do so, we constructed a sinusoidal function of seasonal mite survival within the burrow (Figure 1B, Supplemental Methods), in which the daily probability of mite survival in the simulation varied between a 16-day maximum during the cool-humid winter season and 6-day minimum during the comparatively warmer and dryer summer season. For comparison, we also simulated disease dynamics based on a seasonally invariant burrow environment in which mite survival was stable at 16 days (Supplemental Table 1).

We ran all simulations for 10 years, the estimated average natural life span of healthy wombats (Carver et al. 2024); and set simulations to begin on the first day of winter. The initial conditions of each simulation included a population of 100 wombats with disease entering the system with 10% of individuals having a low severity infection, which is consistent with background prevalence observed in some populations with endemic disease (Driessen et al. 2021, Burgess et al. 2023). Stable and seasonal simulations were each tested over the range of initial ratios of burrows to wombats that have been observed empirically from 1-20 in increments of 0.5 (Martin et al. 2019, Carver et al. 2024).

### 2.2 Characterizing epidemiology with seasonality

#### 2.2.1 Final epidemic states

Over the range of initial burrow to wombat ratios, we compared the final epidemic states of stable and seasonal environments to understand how periodic constraints to mite survival affects long-term persistence of hosts or pathogens. We defined final epidemic states at the end of Year 10 as: host and pathogen extinct, if wombat abundance was less than 1 individual; endemic, if abundance of wombats and abundance of burrows with mites were greater than 1; or parasite extinct, if wombat abundance was greater than 1, prevalence in wombats was 0, and mite colonies were absent from all burrows.

#### 2.2.2 Outbreak dynamics and reservoir persistence over time

To examine the seasonal host and pathogen processes underpinning epidemic outcomes, we focused on scenarios with initial burrow to wombat ratios of 5, 10, and 15 that represent high, moderate, and low relative host-pathogen encounter rates respectively. The selected encounter rates represent previously suggested phenomenon where fewer burrows available per wombat increases the frequency of asynchronous burrow sharing, and thus the rate that a wombat will encounter a burrow with mites present (Beeton et al. 2019). We used the same set of six scenarios (two levels of seasonality and three levels of initial encounter rate, Figure 1B) for all subsequent simulations as they were expected to sufficiently capture seasonal variation in natural contexts. We consider population size (N) to be the absolute number of occupied burrows, local host density (n) to be the fraction of burrows occupied, prevalence (hl) to be the fraction of wombats with low and high severity mange infections, and reservoir persistence (D) as the fraction of burrows that have mites present.

First, to understand temporal variation in host population responses associated with disease, we characterized several epidemiological properties over time including, the frequency, magnitude, and duration of wombat declines and prevalence. Then, we assessed the relationship between host population size and prevalence at each time point, expecting there to be important feedback between these two properties that aligned with the prevalence thresholds for declines previously noted in wombat sarcoptic mange (Carver et al. 2023). Then, we explored differences in overall impacts between stable and seasonal environments for each initial encounter rate scenario by calculating the median, maximum, and minimum prevalence and population size observed over the course of each simulation. We evaluated a similar set of properties in relation to reservoir persistence over time as well as the quantiles of the distribution to describe how seasonality affected variation in the quantity of burrows with mites present in different host contexts. Lastly, we examined how minimizing the temporal disparity between seasonal mite survival and host infection duration influenced disease persistence to understand how the timescales of host and pathogen processes may facilitate co-existence in this system. We simulated the same set of scenarios with shorter time periods for which (i) wombats with mild infection progress to high infection (=15 days) and (ii) highly infected wombats suffer mortality (=30 days) and compared the truncated infectious period outcomes to the natural conditions where infection and mortality are protracted, 30 and 90 days respectively.

The probabilities that a wombat becomes infected from a burrow and a burrow is contaminated by a highly infected wombat are relatively unknown biological parameters. To assess whether these unknown parameters qualitatively changed the population outcome, we ran simulations that increased and decreased each by 10% individually.

### 2.3 Comparison of simulated dynamics to field observations

#### 2.3.1 Data collection

To understand how seasonal forcing of mite survival in the environmental reservoir in the simulations compares to field-based epidemiology, we synthesized data from published field studies that reported location, relative wombat abundance, mange prevalence, and survey date, season or month. We only included studies that used comparable field survey methods (i.e., transect based observations) and were conducted at distinct times and locations. For multiple surveys that occurred within a single season, we calculated the average abundance or prevalence to avoid duplications. When months were reported, we assigned season based on calendar months as winter = June, July, or August; spring = September, October, November; summer = December, January, February; and autumn = March, April, May.

#### 2.3.2 Qualitative analysis

For each simulated scenario and the field dataset, we compared seasonal prevalence trends within a year to our simulated results. We calculated the average prevalence during each quarterly season (i.e., winter, spring, summer, autumn). Then, for each site and year we calculated the relative change between seasons to describe increases or decreases to prevalence during periods when mite survival is expected to be high or low. To account for variation in prevalence at the start of each season, we calculated the proportional prevalence changes as: hl_s+1_ – hl_s_/hl_s_.

## 3 Results

### 3.1 Final epidemic state

Seasonality in mite environmental survival limited the range of local host densities that could support *S. scabiei* persistence in wombat populations relative to stable mite survival (Figure 2). In seasonal environments, endemic outcomes were limited to intermediate burrow:wombats ratios (5-12) whereas high (> 12) and low (< 5) ratios supported disease-free wombat populations with parasite extinction occurring within 4 years (Figure 2, bottom). Stable environments, in contrast, exhibited no upper bound of burrow:wombats ratios that eliminated mites and all burrow:wombat ratios greater than 7 led to endemic disease. Mite extinction occurred at the lowest burrow:wombats ratios in stable environments, but the average time to mite extinction was prolonged (Figure 2, top).

**Figure 2.**
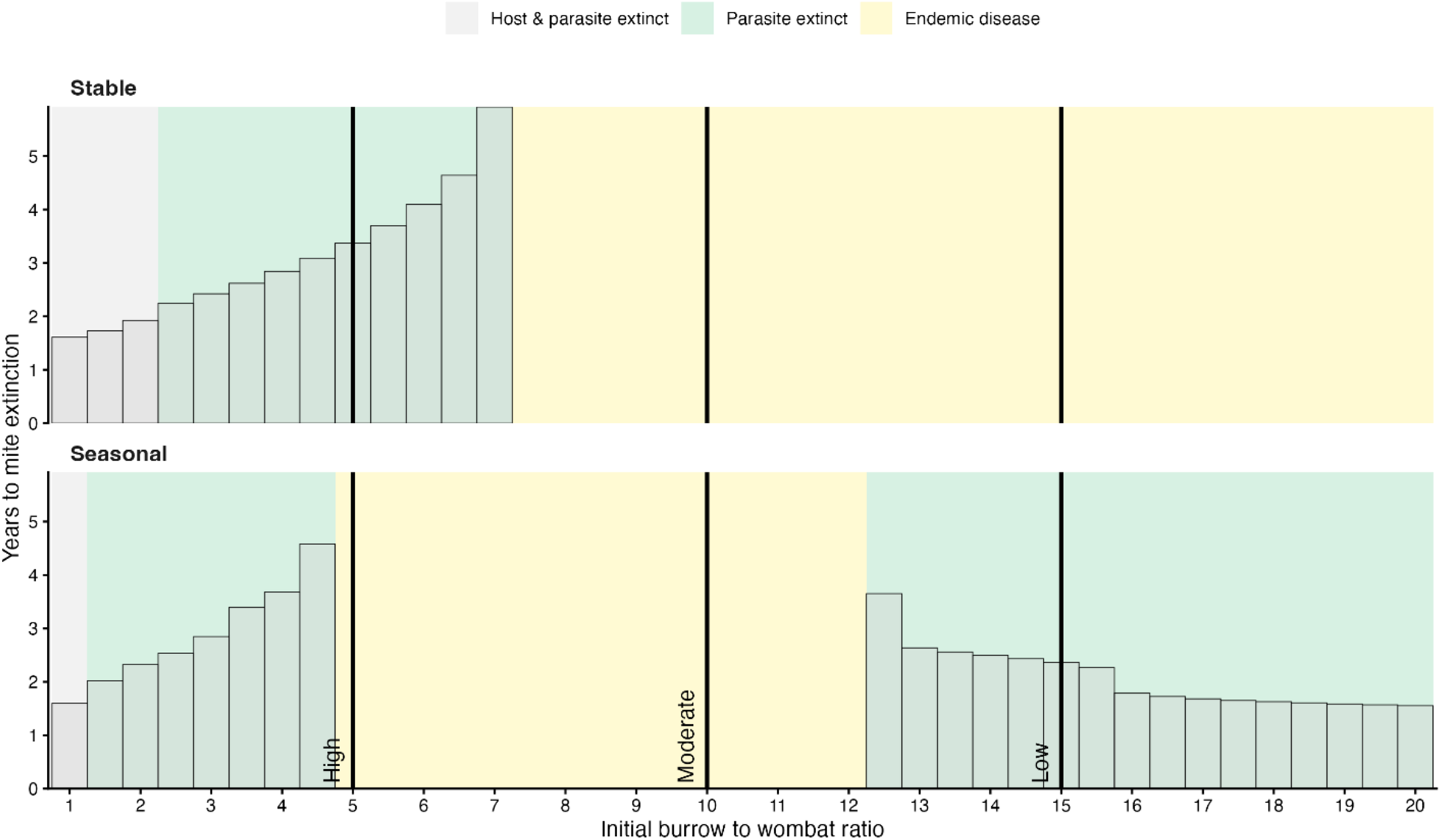
Seasonality in mite survival constrains the range of host-pathogen contexts over which *S. scabiei* can persist in wombat populations. The top and bottom panel show the outcomes in stable and seasonal environments, respectively. Colored rectangles show the epidemic states at the end of the 10-year simulations. The columns show the year when mite colonies went extinct if applicable. Black lines and labels at initial burrow to wombat ratios of 5, 10, and 15 show that initial encounters rates can differentiate long-term trajectories between stable and seasonal environments, which we describe further in subsequent results.

### 3.2 Outbreak dynamics over time

We further explored the epidemiology of burrow:wombat ratios representing three environmental encounter rates – 5 as high, 10 as moderate, and 15 as low – between stable and seasonal environments (Figure 3). In stable environments: high initial host-pathogen encounter rates led to significant temporal variation in host populations (as indicated by the changes in the number of occupied burrows, N) and disease prevalence dynamics (Figure 3A, B); moderate initial host-pathogen encounter rates led to stable host populations and disease prevalence dynamics (Figure 3C, D); and low initial host-pathogen encounter rates produced intermediate host population and disease prevalence dynamics (Figure 3E, F). In contrast, for seasonal environments, intermediate population and disease prevalence dynamics were observed for both high (Figure 3E, F) and moderate (Figure 3C, D) initial host-pathogen encounter rates (similar to low host-pathogen encounter rate scenario for stable environments, Figure 3A, B) and parasite extinction was observed in the low initial host-pathogen encounter rate scenario (Figure 3E, F).

**Figure 3.**
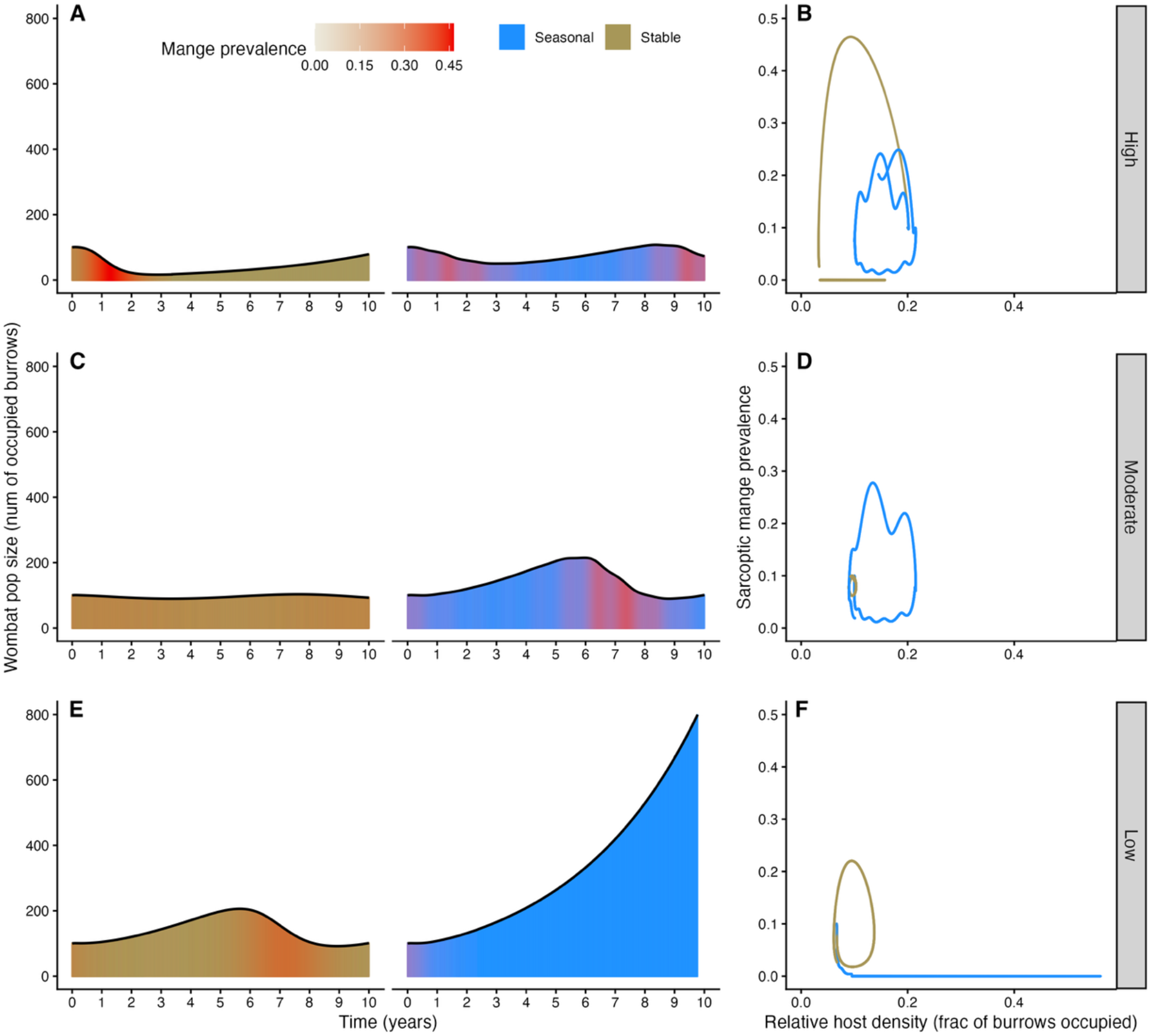
Seasonality in environmental survival of *S. scabiei* varies host outbreaks dynamics with initial host-pathogen encounter rates. Panels A, C, E show temporal variation in population size (black lines) and prevalence (colored columns with red gradient). Panels B, D, F, represent the reciprocal feedback between local host density and prevalence (i.e., outbreak size, hl), illustrating that the relationship between transmission and host density (analogous to encounter rates, burrow:wombat ratios), a key property determining epidemic outcome, is modulated by seasonality.

Examining the temporal dynamics of wombat populations and mange prevalence provided additional insights on how seasonality and host pathogen encounter rates influence wombat-mange dynamics. In the high initial host-pathogen encounter rate scenario, prevalence increased and the wombat population declined followed by recovery in both the stable and seasonal mite survival scenarios (Figure 3A). The degree of decline was less severe in the seasonal mite survival scenario, and the population grew to an extent that a second outbreak and population decline occurred within the 10-year simulation period (Figure 3A, B). In the moderate initial host-pathogen encounter rate scenario, host-pathogen dynamics remained relatively invariant under stable environmental conditions, whereas the population increased leading to an outbreak followed by decline for seasonal mite survival (Figure 3C, D). For the low initial host-pathogen encounter rate scenario, we observed that the population increased leading to a modest sized outbreak and then population decline, whereas parasite extinction occurred with seasonal mite survival leading to exponential population growth (Figure E, F).

We found that the protracted time span of disease progression and mortality is essential for wombat-mite co-existence. After truncating the infectious period from 120 to 45 days (i.e., reducing durations of low to high infection progression from 30 to 15 and highly infected wombat survival from 90 to 30 days) to be similar to the daily changes in mite survival, parasite extinction was rapid and host populations were released from disease in all scenarios (Supplemental Figure 1). The population trajectories that resulted from increasing and decreasing the relatively unknown parameters produced qualitatively similar differences in epidemiology between stable and seasonal environments (Supplemental Figures 2-3), such that seasonal environments limit endemic outcomes and generally supported higher population sizes.

### 3.3 Seasonal effects on environmental reservoir abundance

Seasonality varied the accumulation of contaminated reservoirs (fraction of burrows with mites present) over time across the three initial host encounter rates (burrow to wombat ratios of 5:1, 10:1, and 15:1; Figure 4A, B). In stable environments: high initial encounter rates led to an early peak in the reservoir that was the greatest in magnitude among all scenarios (Figure 4A, top), intermediate initial encounter rates had the least extensive reservoir accumulate over time (Figure 4A, middle) and was also the least variable (Figure 4B, middle), and low initial encounter rates delayed reservoir establishment until the latter half of the simulation and was maintained only to a moderate extent (Figure 4A, bottom).

**Figure 4.**
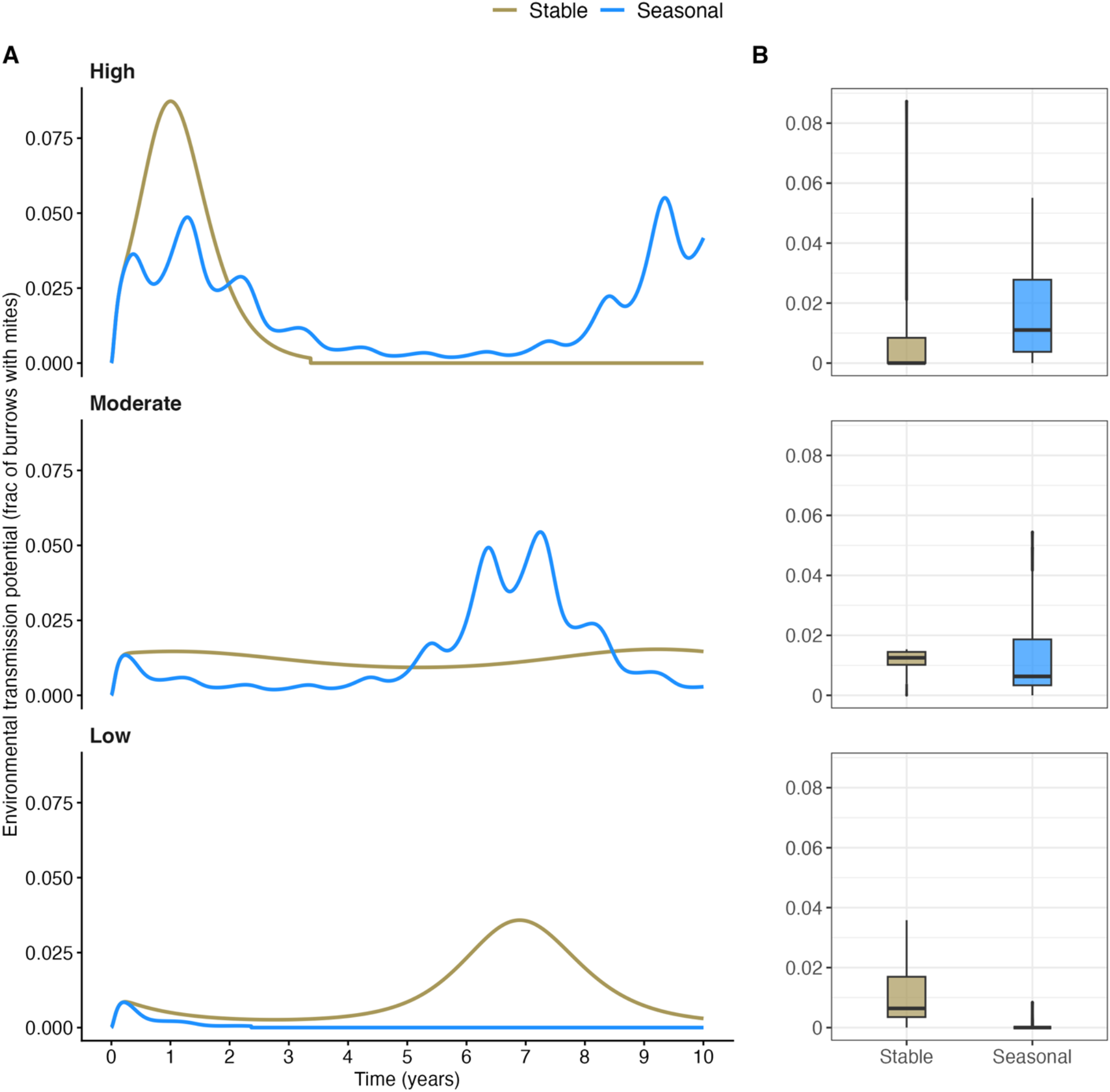
Temporal variation in environmental reservoir abundance, as the fraction of burrows with mites present, under high (top panels), moderate (middle panels), and low (bottom panels) initial host encounter rates. Tan and blue colors show output from stable and seasonal scenarios, respectively. **A)** The fraction of burrows with mites present at each time step showing that oscillations of the reservoir in the seasonal environment are synchronized with mite survival but vary in amplitude. **B)** Summarized variation over the course of the simulation by initial host encounter rate that corresponds with the encounter rates of Panel A.

In seasonal environments: the contaminated reservoirs accumulated quickly in all initial encounter rates (Figure 4A). As the epidemic progressed in seasonal environments, high initial encounter rates supported two periods of reservoir accumulation, or peaks of environmental fomites (Figure 4A, top), consistent with population and prevalence patterns in the host (Figure 3). For intermediate encounter rates, the environmental reservoir did not accumulate significantly until later in the simulation (Figure 4A, middle), and low initial encounter rates inhibited reservoir establishment (Figure 4A, bottom).

### 3.4 Comparison of simulated dynamics to field observations

The output from our simulated models were generally consistent with field observed prevalence trends and population declines. We found that the seasonal simulations produced prevalence patterns more like field data than stable simulations (Figure 5), such that a prevalence increase was more apparent during seasons when the environmental conditions are more suitable to longer mite survival in the environment and was less apparent when the environmental conditions are less suitable to mite survival in the environment. As expected, there was no difference in mange prevalence over time in the stable dataset (Figure 5). We also determined there were not site or survey year biases that contributed to the empirical trends (Supplemental Figure 4).

**Figure 5.**
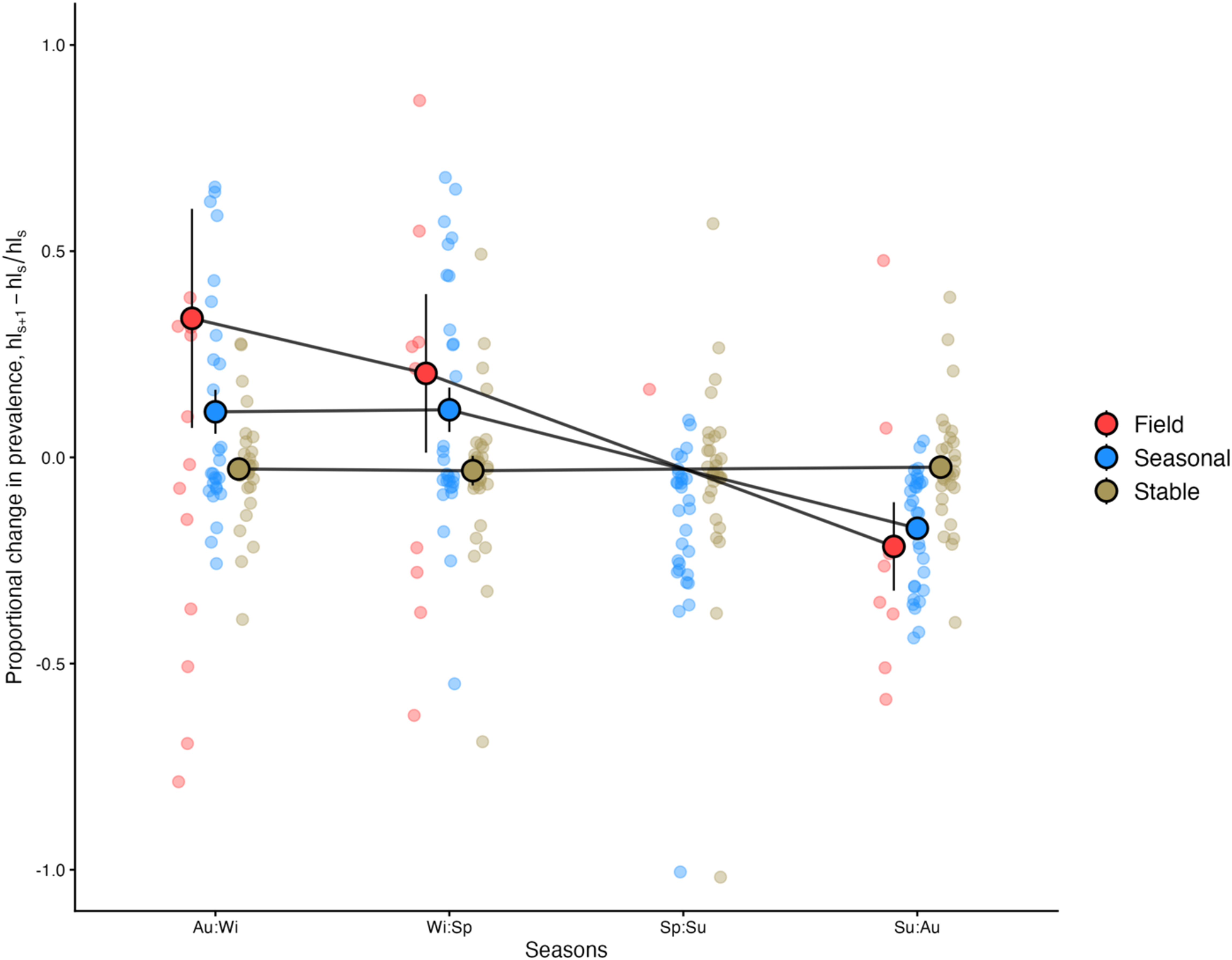
Differences in the relative change in prevalence change between seasons. Small points represent a season-to-season change in prevalence within a year at a distinct survey location. Points with the black outlines and vertical bars represent the mean and standard errors. Spring to summer prevalence change was excluded due to a lack of field data reporting consecutive prevalence in a single location. Calculations for the proportional change in prevalence (hl) were based on s, as the initial season, and s+1, as the subsequent season.

## 4 Discussion

Seasonal effects on pathogen survival in abiotic reservoirs are fundamental to the epidemiology of environmentally transmitted pathogens, yet when the timescale of infection and disease in the host are decoupled from the parasite in the environment, the dynamical impacts of this are poorly understood in wildlife systems. Here, we used a theoretical framework informed by field and lab measurements to better understand variation in wombat population responses to sarcoptic mange, where the parasite has seasonal environmental survival and the host experiences a protracted clinical infectious period. We demonstrated three different epidemic outcomes that were controlled by seasonality in mite environmental survival: host and parasite extinction, parasite only extinction (disease-free), and endemic disease. Interestingly, seasonal constraints on mite survival did not consistently reduce the presence of mites in burrows to minimize outbreaks or declines. Rather, seasonal mite survival interacted with initial host encounter rate to mediate epidemics, indicating that the effect of seasonality on long-term outcomes differs with hosts contexts, and seasonal controls on epidemics may not be apparent given the disparate time scales in which host and pathogen demography are altered by seasonal conditions.

Identifying the factors that control population responses to invading pathogens is key to evaluating the consequences of disease and prioritizing management. We found that seasonal constraints on mite survival due to fluctuations in temperature and humidity governed the epidemic outcome of wombat populations affected by sarcoptic mange. We observed three possible outcomes and determined that the range of host conditions that support endemic disease is narrowed by seasonally forced reservoir persistence. Each of the epidemiological states we projected are also observed in empirical field studies. Near extinction of wombats have been observed at Narawntapu National Park with a decline of 94% in the wombat population over 7 years, while endemic disease occurs at Cape Portland in north-eastern Tasmania and New South Wales (Martin et al. 2018, Stannard et al. 2020, Driessen et al. 2021). Disease extinction and total extinction outcomes are harder to quantify, but wombat populations exist free of mange, despite the parasite being in Australia for >200 years, suggesting parasite extinction has possibly been common (Driessen et al. 2021). The epidemic outcomes we described are also consistent with three of four outcomes predicted by the SIR framework from Beeton et al (2019), which additionally observed stable limit cycles at low population abundance, which may or may not occur in this system naturally. The addition of seasonality to our model appears to deduce populations outcomes that align more closely with empirical observations than the former model which lacked a mite survival parameter that fluctuated seasonally.

Environmental reservoirs, and reducing their persistence on the landscape, are a common target of disease control to minimize transmission potential (Langwig et al. 2015). Counterintuitively, periodic reductions in mite survival, explored here as natural seasonal fluctuations, did not consistently benefit long-term population outcomes. Our results indicate that wombat populations are released from disease under two general circumstances: an outbreak is severe enough to reduce populations to sizes insufficient for transmission or a high availability of burrows minimizes burrow sharing and reduces transmission. We highlight the importance of considering host contexts when deciding whether to focus on environmental reservoir management. We also found that seasonal reductions in mites supported higher population sizes generally, so in locations with endemic disease or if endemism appears inevitable, reservoir management may be a useful strategy if the goal is to maintain larger host populations. While beyond the scope of this study, future management actions can expand this model using a Model Integrated Disease Management approach similar to what has been applied to this system formerly and others (Carver et al. 2022, Grant et al. 2024, Grogan et al. 2025), to evaluate other regimes through which mite survival can be effectively reduced to support wombat populations, such as increasing the frequency or efficacy (i.e., magnitude) of interventions. For example, reducing mite persistence through burrow microclimate manipulations or environmental treatments or altering the availability of burrows by creating artificial habitats could be valuable avenues.

Periodicity of transmission and host population sizes over different timescales may generate inapparent seasonal effects that are easily overlooked. Mite survival matches temporal fluctuations in burrow climatic conditions with a decline in summer and peak in winter (Browne et al. 2021). Our model, consistent with field observations, shows host population fluctuations and annual growth rates are influenced by climate cycles through periodic transmission, but are asynchronous with off-host mite survival fluctuations (Martin et al. 2018, Stannard et al. 2020, Driessen et al. 2021, Carver et al. 2023). There was a serial lag between seasonal mite reservoir reduction, prevalence change, and wombat population change. Once the reservoir established, the persistence of mite-present burrows oscillated in concert with seasonal climate cycles, however, the amplitude of the peaks varied over time (i.e., how much it increased or declined seasonally). These lag effects as well as reservoir maintenance are likely reflecting the relatively slow life history of wombats, with maternal reproductive cycles ca.1.5 years apart, an estimated natural 10-year lifespan, and a slower disease progression compared to the rate that mite survival changes (Martin et al. 2018, Martin et al. 2019, Browne et al. 2021, Yabsley et al. 2025). The slow life history of wombats may also contribute to why wombats have not yet evolved better defenses to *S. scabiei* and how mites are able to persist through seasonal, climate-induced population bottlenecks. Long gestation, extended parental care and relatively slow rates of recruitment potentially slow the evolution of host resistance (Valenzuela-Sánchez et al. 2021) and, being coupled with relatively slow disease progression that extends transmission opportunities, may be ameliorating selection on the pathogen to reduce virulence. The strongest selection pressure on mites may come from burrow microclimates and *S. scabiei* may reach an endemic state in wombats by optimizing off-host survival as predicted for environmentally persistent pathogens (Walther et al. 2004).

A lack of strong seasonal behaviors and physiology or direct contact transmission could dampen, rather than exacerbate, seasonal infection dynamics that are evident in other host species. Ibex, foxes and humans, for instance, have predictable annual prevalence peaks for *S. scabiei* that is attributed to elevated seasonal contact rates among individuals (López-Olvera et al. 2015, Azene et al. 2020, Iacopelli et al. 2020, Loredo et al. 2020). In contrast to the other mange-affected species where seasonal traits of hosts drive clear, relatively synchronous prevalence peaks, the contribution of seasonality in wombat-mange dynamics is less apparent because it operates indirectly via the reservoir but plays out at longer time scales for the host. Future work aiming to understand the long-term disease outcomes in other solitary species impacted by sarcoptic mange, such as the recent emergence of sarcoptic mange in American black bears (Niedringhaus et al. 2019, Niedringhaus et al. 2019, Francisco et al. 2025), should assess seasonal factors expected to periodically affect transmission.

Population regulation between hosts and pathogens is a key principle in disease ecology (Anderson et al. 1979, May et al. 1979) but is difficult to identify in natural systems. We observed reciprocal feedback between relative host density and prevalence in the wombat-mange system that was mediated by seasonality and initial host population size. Field data on New South Wales wombat populations show that the apparent prevalence of mange ranges from 7% up to 41% (Stannard et al. 2020), and in Tasmania was observed up to 32% in populations surveyed and varied spatially (DPIPWE 2020), broadly consistent with our model predictions that indicated prevalence does not exceed 37%. Likewise, burrow occupancy, or local population density, within the model, does not exceed 21.5% across all scenarios equating to a 4.7:1 burrows:wombat, a relatively high encounter rate. Data on burrow:wombat ratios is limited and not all are in populations with mange present. Surveys in Tasmania from mange infected populations suggest burrow occupancy has been observed at 3.5:1, 4.3:1 and 11.1:1 burrows:wombats (Roger et al. 2007, Martin et al. 2019, Burgess et al. 2023). Populations of wombats can reach much lower burrows:wombats ratios without the limitation *S. scabiei* imposes on burrow occupancy, ranging from 11:1, 3:1 to as low as 1.1:1 in high density pastoral land with no mange present (Skerratt et al. 2004, Carver et al. 2024). By synthesizing the data on field studies to understand relationships in declines and the prevalence of infection, we found that the qualitative relationship between an annual decline in simulated environments was like field-based patterns, such that annual declines are expected to occur when prevalence is >10%. Cumulative increases in prevalence between years could explain long-term population declines in other studies (Carver et al. 2023), and alternatively, periodic reductions in the reservoir due to seasonality may hold prevalence at or below this threshold, allowing populations to be stable.

While we replicated many aspects of *S. scabiei*-wombat dynamics observed in nature, we also highlight several areas of future study that would minimize discrepancies between our model output and sarcoptic mange in wombats in natural systems. Firstly, burrow to wombat ratio in our model is only altered through birth and death rates of the wombat population with burrow abundance remaining constant, but the number of burrows within a population likely changes over time due to burrow abandonment, the creation of new burrows, or temporary inactivity of existing burrows (Evans 2008). Secondly, the model assumes random burrow selection by wombats, but studies suggest landscape factors (such as local habitat) influence preferential burrow use (Burgess et al. 2023) or more nuanced movement dynamics between burrows should influence transmission, as noted in other systems with spatially distributed reservoirs (George et al. 2013, Turner et al. 2013, Leach et al. 2016, Langwig et al. 2021, Dolfi et al. 2024). Quantifying both aspects of burrow use in the field would enhance model realism by providing more accurate estimates of indirect transmission. Lastly, the environmental conditions we used to parameterize seasonal mite survival and describe field declines were primarily from coastal regions of Australia. Future studies that measure microclimates and relative burrow to wombat ratios for inland populations would encompass a broader climate gradient and allow for a more comprehensive assessment of disease dynamics.

Seasonal forcing of epidemics is difficult to characterize for diseases with environmental transmission, particularly when the infectious period and demography of the host, is relatively slow compared to parasite persistence rates. Here, we demonstrate the benefit of integrating field and lab investigations into a model framework to better understand empirically observed population outcomes of an important species facing disease challenges. Scaling seasonal processes to population outcomes is essential for forecasting the persistence of invading pathogens, but we demonstrate that seasonality may be both a promotor and inhibitor of endemic disease under different host conditions.

## Supporting information

Suppemental methods, figures, and table

## Author contributions

M.J.K.: conceptualization, data curation, formal analysis, methodology, visualization, writing - original draft, writing - review & editing, L.C.: conceptualization, formal analysis, methodology, visualization, writing – original draft, M.V.: conceptualization, writing – review & editing, S.A.R.: conceptualization, formal analysis, methodology, supervision, writing – review & editing, S.C.: conceptualization, formal analysis, methodology, supervision, visualization, writing – original draft, writing – review & editing.

All authors gave final approval for publication and agreed to be held accountable for the work performed therein.

## Acknowledgements

We thank the Australian Research Council Linkage Project (LP180101251) for financial support to Scott Carver and Shane A. Richards. We also thank members of the Center for the Ecology of Infectious Diseases at University of Georgia for insightful discussions and the researchers whose previous work contributed to constructing and evaluating the mathematical model.

## Statement on inclusion

Our study was a collaboration among researchers from multiple countries, including scientists based in the study region who engage with local stakeholders and practitioners. All authors’ contributions were valued throughout project, from the onset of study design and through later stages of drafting, to ensure diverse perspectives were reflected in the content. All the relevant works, including model framework and field data, that were synthesized from the literature have been thoroughly cited and acknowledged.

## Data availability and sources

The code and data used in this study will be made publicly available at Dryad Digital Repository.

