## Supplementary material for "Decoupled seasonal effects of an environmentally transmitted wildlife disease": Suppemental methods, figures, and table

**Supplemental material for Decoupled seasonal effects of an environmentally transmitted wildlife disease**

### Methods

*Expanded model description*

a) System overview

As in Martin et al. 2019, our model represents an environment that is comprised of many burrows, each belonging in one of eight states that can change daily. The maximum number of wombats that occupy a burrow is equal to 1. Burrows become contaminated when mites are transferred from infected wombats and wombats acquire infection when mites are transferred from a contaminated burrow (burrows with mites present).

On day $t$, a randomly selected burrow has the probability of being in each state as $P_{i,j,t}$, based on host occupancy status, $i$ = {U, S, L, H}, and reservoir status, $j$ = {D, d}. Host occupancy status is as follows: U = unoccupied, S = occupied by susceptible wombat, L = occupied by wombat with low infection, or H = occupied by wombat with high infection. Reservoir status is D = contaminated, mites present or d = mites absent.

Per burrow wombat population density on day $t$ is equal to the number of occupied burrows as:

$$N_t = \sum_{i \neq \mathrm{U}, j} P_{i,j,t}$$

Contaminated reservoir density on day $t$ is given by the fraction of burrows that contain mites, either persisting off-hosts or being occupied by an infected wombat as:

$$D_t = \sum_{i, j = \mathrm{D}} P_{i,j,t}$$

b) State dynamics

The state dynamics of our system are generally consistent with the state dynamics described by Martin et al 2019, however, we excluded the daily events specific to their treatment objective (i.e., treated wombats lose immunity, treated burrows lose treatment viability, and burrows gain effective treatment) and the *k* parameter representing treatment status. The pre-existing model structure was developed flexibly to exclude treatment states and the associated event transition probabilities by setting the number of treated burrows to 0. As a result, our state dynamics mirror the set of equations for wombat deaths, disease persistence in burrows, wombat disease progression, wombat burrow switching, and wombat gains independence, with the wombat occupancy states associated with immunity from treatment ($i = I$) or burrows treatment status (k = {T, t}) are muted mathematically.

Accordingly, $P_{i,j,t}^{(i)}$ denotes the burrow states on day $t$ after event $i$, so the distribution of burrow states within the given environment at the beginning of each day can be calculated by $P_{i,j,t+1} = P_{i,j,t}^{(5)}$. See Appendix S4.3 of Martin et al 2019 for the full sets of equations used for the state dynamics. Our extension of the model to include seasonal forcing in burrow reservoirs is achieved through the disease persistence in burrows event probability (Appendix S4.3.2 of Martin et al 2019), where $f$ is the number of days it takes for a contaminated burrow (i.e., associated with a diseased host or mite presence) to lose mites. We transformed $f$ to a time-varying value, $f\_t$, such that the daily probability a contaminated burrow becomes mite-free then also becomes time-dependent as $P_{d\_t} = 1/f\_t$.

Using a sinusoidal function to reflect seasonal fluctuations built with empirical estimates of temperature and humidity dependent mite survival off hosts, we find $f\_t$ for each day of the simulation based on:

$$f_{min} + 0.5(f_{max} - f_{min}) * (1.0 + \cos\left(2\pi * \frac{T}{365}\right))$$

where $f_{min}$ equals the lowest mite survival time in days, $f_{max}$ equals the highest mite survival time in days, and $T$ is the duration of the simulation scaled to biological seasons.

**Figures**

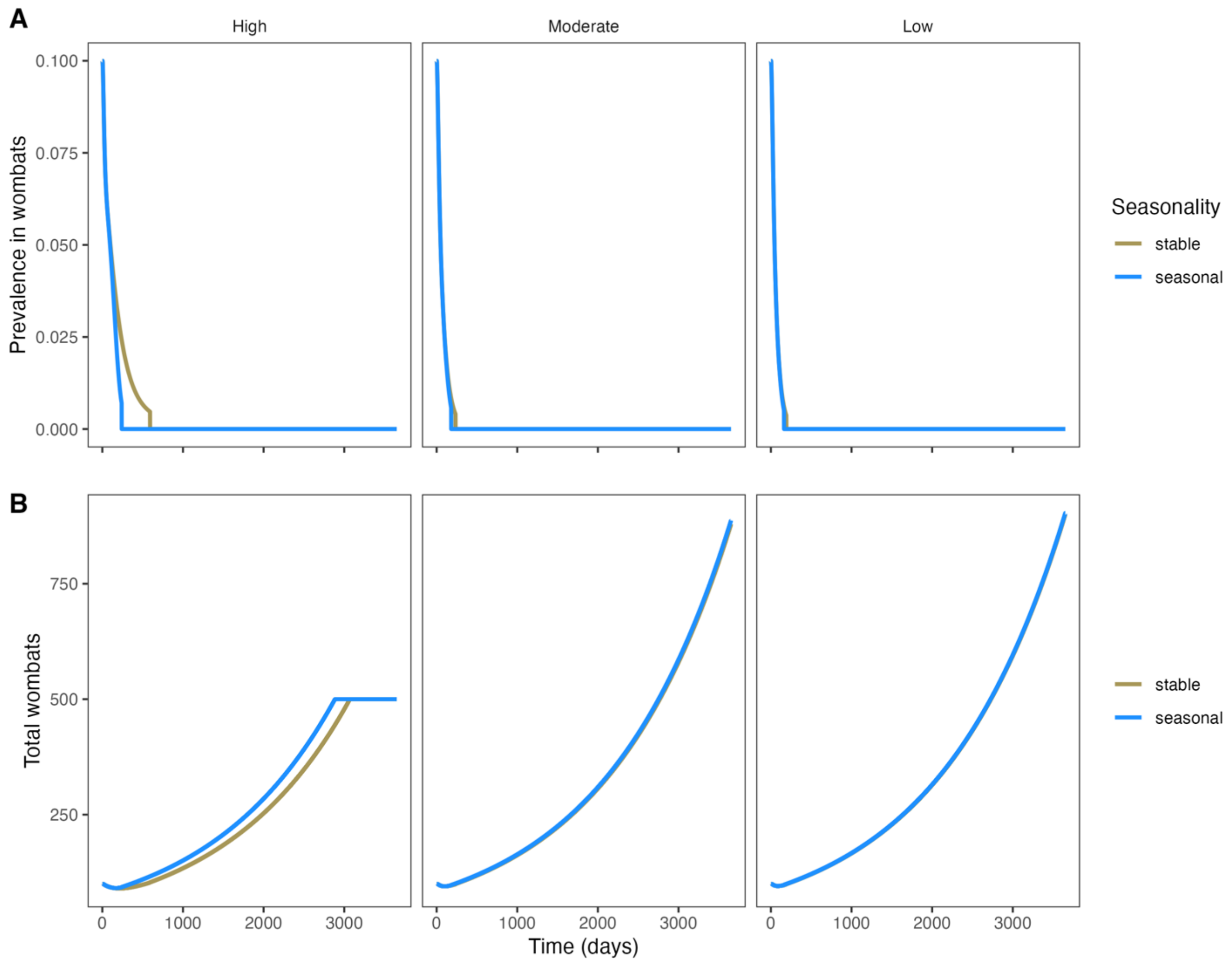


**Supplemental Figure 1.** Wombat-mange fails to establish when the infectious period is truncated (low to high infection progression from 30 to 15 days; high infection to mortality from 90 to 30 days). Minimizing the temporal disparity between seasonal parasite survival, infection and disease-associated mortality inhibited parasite persistence **(A)** and released wombat populations from disease **(B)** in all scenarios. Panels from left to right show the relative host encounter rates based on wombat:burrow ratios 5, 10, and 15 respectively. Line colors denote stable or seasonal environments.

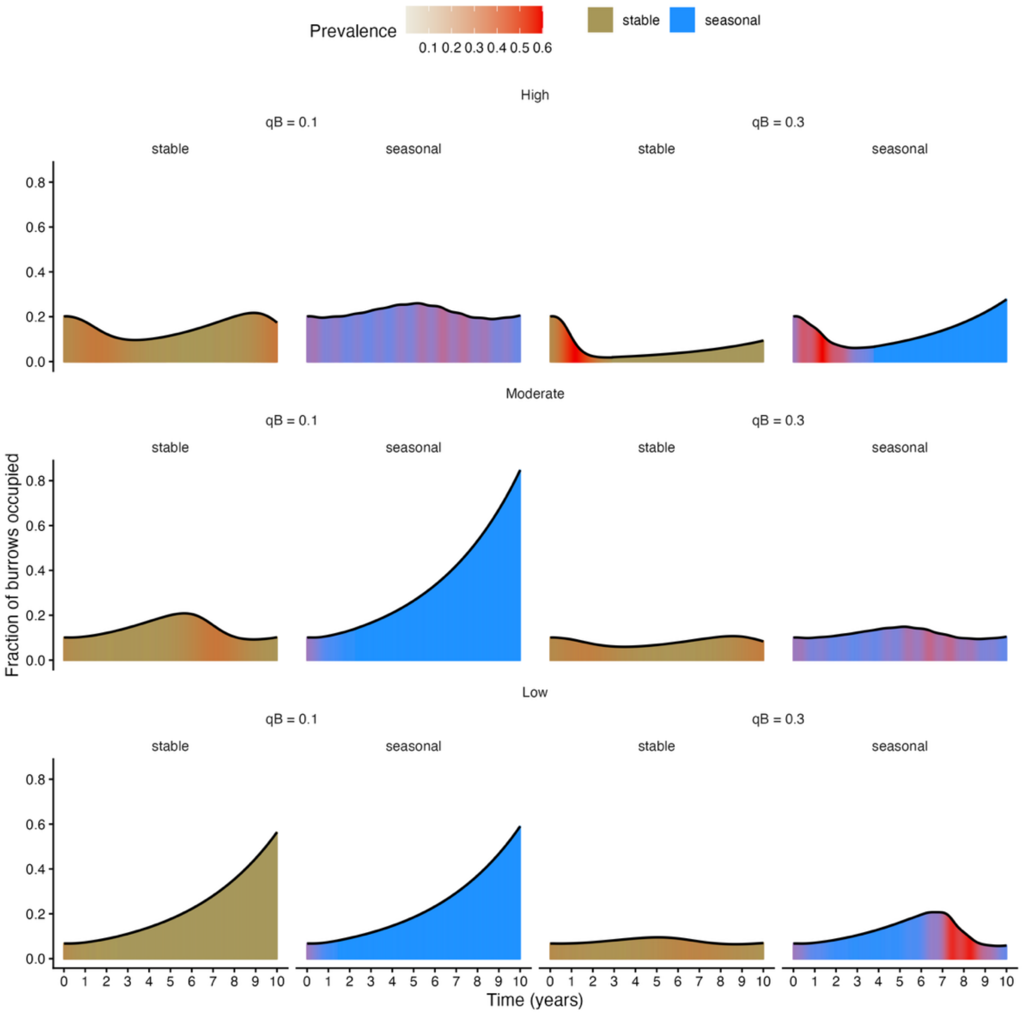


**Supplemental Figure 2**. Temporal dynamics of relative host density and prevalence in wombats when the probability of acquiring infection from a burrow, qB, is decreased (left panels) or increased (right panels) by 10%. Rows from top to bottom show decreasing relative initial encounter rate as high, moderate, and low, respectively.

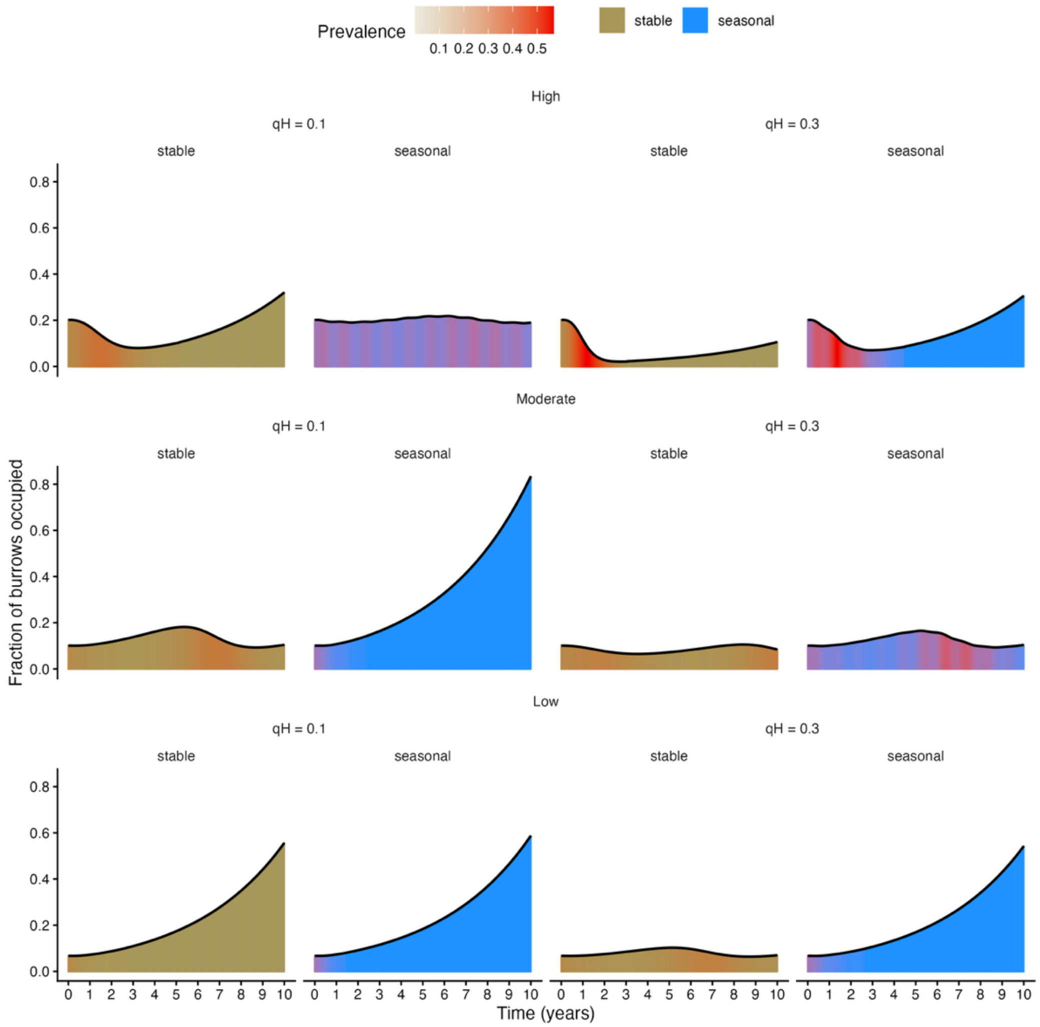


**Supplemental Figure 3**. Temporal dynamics of relative host density and prevalence in wombats when the probability of a highly infected wombat contaminating a burrow is decreased (left panel) or increased (right panel) by 10%. Rows from top to bottom follow decreasing relative encounter rates as high, moderate, and low, respectively.

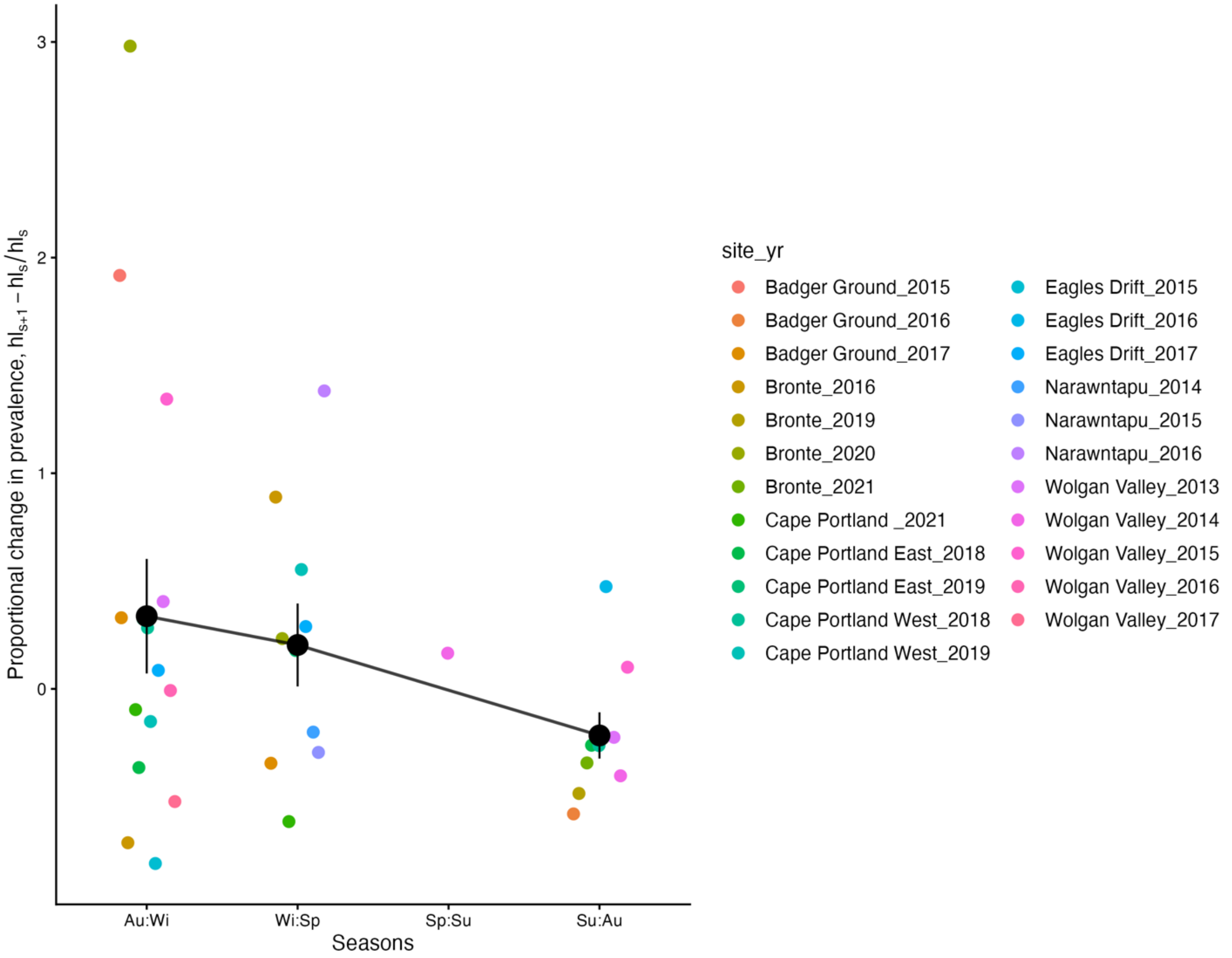


**Supplemental Figure 4.** Relationship between the proportional change in prevalence between consecutively surveyed seasons (s to s+1) based on the synthesized field data. Points are colored by the site and year of survey. Black points and vertical bars show the average across survey with a line showing the relative trend. No survey site or year obviously biased the prevalence trends, whereby increases in prevalence primarily occurred during seasons when mite survival also increased (i.e., autumn to winter and winter to spring).

### Tables

**Supplemental Table 1.** Relevant parameters used within the state-based, discrete time model for all scenarios. Model symbols correlate to supporting information. All parameters were based on Martin et al 2019 except for seasonal mite survival that was estimated in Browne et al 2021.

| Model Parameter | Model Value | Model Symbol |
|---|---|---|
| Wombat birth rate (1 joey/1.5 years) | 547.5 days | Tau |
| Wombat life expectancy (10 years) | 10*365 days | L |
| Wombat life expectancy at low infection | 10*365 days | LL |

| | | |
|---|---|---|
| Wombat life expectancy at high infection | 90 days | LH |
| Low-high infection duration | 30 days | D |
| Burrow switching frequency | 0.13/day | P |
| Mite survival, stable environment | 16-16 days | F(min) – F(max) |
| Mite survival, seasonal environment | 6-16 days | F(min) – F(max) |
| Prob. An infected burrow infects a wombat | 0.2 | qB |
| Prob. A highly infected wombat infects a burrow | 0.2 | qH |
| Initial burrow to wombat ratio as encounter rate (High, Moderate, or Low) | 5, 10, or 15 | |
